# Microstimulation reveals anesthetic state-dependent effective connectivity of neurons in cerebral cortex

**DOI:** 10.1101/2024.04.29.591664

**Authors:** Anthony G Hudetz

## Abstract

Complex neuronal interactions underlie cortical information processing that can be compromised in altered states of consciousness. Here intracortical microstimulation was applied to investigate the state-dependent effective connectivity of neurons in rat visual cortex in vivo. Extracellular activity was recorded at 32 sites in layers 5/6 while stimulating with charge-balanced discrete pulses at each electrode in random order. The same stimulation pattern was applied at three levels of anesthesia with desflurane and in wakefulness. Spikes were sorted and classified by their waveform features as putative excitatory and inhibitory neurons. Microstimulation caused early (<10ms) increase followed by prolonged (11-100ms) decrease in spiking of all neurons throughout the electrode array. The early response of excitatory but not inhibitory neurons decayed rapidly with distance from the stimulation site over 1mm. Effective connectivity of neurons with significant stimulus response was dense in wakefulness and sparse under anesthesia. Network motifs were identified in graphs of effective connectivity constructed from monosynaptic cross-correlograms. The number of motifs, especially those of higher order, increased rapidly as the anesthesia was withdrawn indicating a substantial increase in network connectivity as the animals woke up. The results illuminate the impact of anesthesia on functional integrity of local circuits affecting the state of consciousness.

## Introduction

Neuronal connectivity underlies information processing in the brain. Exogenous perturbation of discrete elements of neuronal circuits can help reveal the effective connectivity of neurons beyond what can be inferred from recording of undisturbed, spontaneous activity (Clark et al., 2011;Kumar et al., 2013;Sadeh and Clopath, 2020). The effect of anatomically precise local perturbations, even a single extra spike, can propagate throughout the neuron network, generating multiple extra spikes that last for hundreds of ms (London et al., 2010;Kwan and Dan, 2012;Emiliani et al., 2015;Bernardi et al., 2021). Such spike sequences formed by forward and recurrent interactions may reflect basic information packets in the neuron network (Luczak et al., 2015).

Recurring spike patterns are also state dependent; they are affected by various physiological and pharmacological manipulations. A recent study demonstrated that different levels of anesthesia produce specific changes in spike sequences in visual cortex (Tanabe et al., 2023). Such changes in spike patterns may inform the mechanism by which state-dependent cellular changes could impair critical elements of neuronal information processing and whose breakdown ultimately leads to loss of consciousness as produced by anesthetics and other interventions (Hudetz et al., 2015;Margalit et al., 2022;Pazienti et al., 2022).

Prior works emphasized the potential of intracortical microstimulation to probe the structure and function of neuronal circuits (Clark et al., 2011;Kwan and Dan, 2012;Kumar et al., 2013). However, the systematic effect of anesthesia on microstimulation-induced changes in effective connectivity of local circuits has not been reported. Anesthetic modulation of neuronal connectivity has been traditionally derived from recordings of spontaneous ongoing activity or sensory stimulation-evoked activity (Hudetz et al., 2009;Andrada et al., 2012;Vizuete et al., 2012;Aasebo et al., 2017;Lee et al., 2021;Bharioke et al., 2022;Margalit et al., 2022;Aggarwal et al., 2023). Recently, intracortical stimulation was employed to examine the effect of anesthesia on electroencephalographic connectivity (Arena et al., 2021) but not yet in spiking neuronal networks. Here intracortical microstimulation was employed to investigate how anesthesia at graded levels may alter intracortical stimulation-related spiking activity and effective connectivity (Aertsen et al., 1989) of discrete neurons *in vivo*. Specifically, the study aimed to determine how microstimulation at a single electrode site alters the spiking activity of neurons at other sites of the same electrode array, repeated for all available stimulation sites, thus sampling the effective connectivity of the underlying neuron network. This work yields insight into how anesthesia impacts the functional integrity of local cortical circuits.

## Materials and Methods

### Surgery and Experimental protocol

The study was approved by the Institutional Animal Care and Use Committee in accordance with the Guide for the Care and Use of Laboratory Animals of the Governing Board of the National Research Council (National Academy Press, Washington, D.C., 2011). General procedures followed those described before (Lee et al., 2020) but using a different probe design. Briefly, nine adult male and female rats were chronically implanted with microelectrode arrays (Microprobes, Gaithersburg, MD, USA) into the right primary visual cortex (V1) for extracellular recording and stimulation. The probes consisted of 32 platinum/iridium (70%/30%) electrodes of 75 μm diameter and 2mm length. They had approximately 100 kOhm impedance. The electrodes were arranged in a 6x6 square matrix format, omitting the corner locations. The electrode spacing was 250 μm in both directions. The stereotaxic target coordinates of the center of the array at the electrode tips were -6.75 mm anterioposterior, 3.60 mm mediolateral, and 1.50 mm deep from Bregma. The depth was chosen to target infragranular cortex (layers 5/6); this was not verified with histological analysis in this study. One corner of the electrode array was occupied by the reference electrode consisting of the same Pt/Ir composition with 10 kOhm impedance. A bare wire tied to a stainless-steel screw in the cranium over the left cerebellum served as ground.

Several days after recovery the animals were placed unrestrained in a sealed, dark anesthesia chamber for testing. The experiment consisted of stepwise decrease of inhaled concentration of desflurane at 6%, 4%, 2.5% and 0% mixed to 30% oxygen-enriched air. Desflurane concentrations were chosen to cover a suitable range from light sedation to deep anesthesia. Based on our former experiments (Imas et al., 2005) with testing the righting reflex - a putative index of consciousness in rodents (Franks, 2008), 2.5% corresponds to conscious sedation, 6% unconsciousness and 4% is near the transition point. Anesthetic depth was not tested in the present experiments. Each anesthetic level was held for 60 minutes including an equilibration time of 15 minutes before commencing microstimulation. A typical experiment lasted for 4 hours.

### Stimulation parameters

Monopolar stimuli were delivered to each electrode site one by one in random order at 0.5s intervals (2Hz). The stimulation frequency was chosen to maximize the number of stimuli delivered while allowing population spike activity return to the prestimulus baseline. The same stimulation sequence was repeated 160 times, equivalent to a total of 2560 stimuli delivered in each condition. Stimulation pulses were charge-balanced, biphasic waveforms (Merrill et al., 2005) that were initially varied with respect to cathodic-anodic temporal order and asymmetry of duration and amplitude. Preliminary tests with 5, 10, 20, and 40 μA stimulation currents showed that maximum response was afforded by 40 μA in agreement with other studies (Voigt and Kral, 2019;Sombeck et al., 2022;Yun et al., 2023). Therefore, in subsequent experiments a maximum stimulus current of 40 μA was used. The amplitudes and phase durations for asymmetric biphasic stimuli were 40 μA for 60 μs, 0 μA for 60 μs, and 8 μA for 300 μs.

### Recording and Data analysis

Extracellular potentials were recorded at all electrode sites except at the one being stimulated using the data acquisition system, Scout Processor and Nano2+Stim front end (Ripple Neuro, Salt Lake City, UT, USA), digitized at 30kHz and band-pass filtered at 300-7500 Hz for spike detection. Unit spikes were extracted and sorted using the software Spyking Circus (Yger et al., 2018). The stimulus artefact was removed by template subtraction (Sombeck et al., 2022). The number of active neurons and those responding to microstimulation varied from trial to trial. A total of 160 spiking neurons were recorded in 9 rats. Neurons were classified off-line into putative excitatory or inhibitory type based on the half peak-width and trough-to-peak time of their spike waveform as illustrated in **Figure 1A** and **Figure 1B**. (Lee et al., 2020). **Figure 1C** shows original recordings of an excitatory and an inhibitory neuron as an example. Effective connectivity maps were constructed from all pairs of stimulus-neuron sites from neurons with a significant change in spike rate within 1-4 ms after stimulation. Spike counts from 20 trials at each stimulus site were averaged. Significant change was defined by a spike rate increase (or decrease) exceeding ±95% confidence interval of the 200 ms pre-stimulus baseline. Putative monosynaptic connections were determined from short time lag spike cross-correlograms (CCGs) from spontaneous and stimulation-induced activity as before (Vizuete et al., 2012;Kobayashi et al., 2019). Briefly, for each pair of neurons, CCGs were calculated at 1 ms time bins over an interval of ±50 ms. To avoid false detection due to background fluctuations, the CCGs were also calculated from surrogate spike trains prepared by “jittering” the spike times within ±5ms, repeated 1000 times to yield 1000 surrogate data sets. The 99% confidence interval of the number of CCG counts in each time bin were obtained and their maximum and minimum were used as global thresholds. Putative excitatory or inhibitory monosynaptic connections between a pair of neurons were inferred by the presence of CCG peak exceeding (excitatory) or trough descending below (inhibitory) the respective global threshold within 1-4 ms time lag (**Figure 1D**). Within-group effects of microstimulation and anesthetic level were tested with ANOVA followed by Tukey Honestly Significant Difference post-hoc tests.

**Figure 1.**
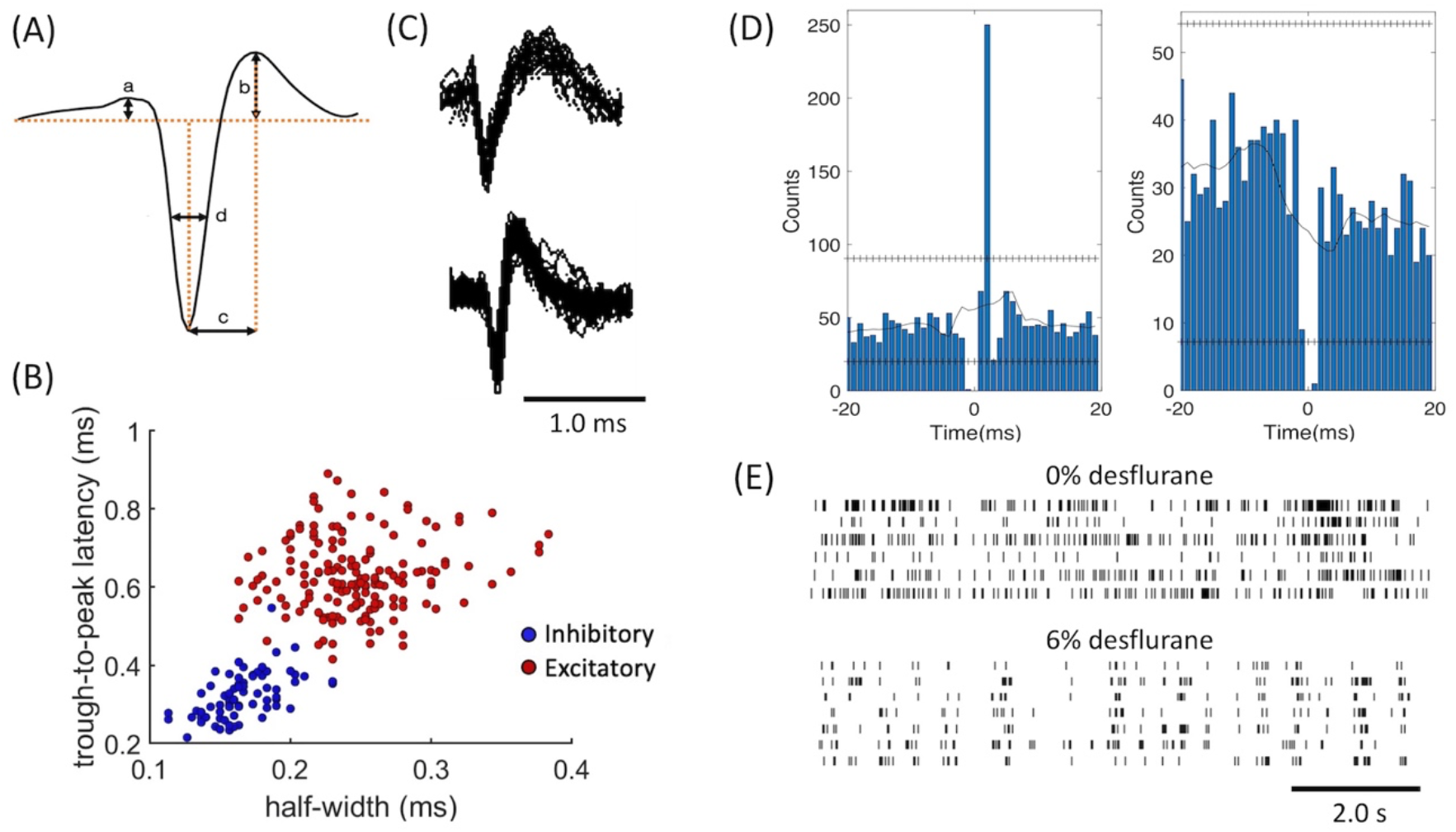
Classification of units and monosynaptic connectivity. **(A)** Features of spike waveform used for unit classification. **(B)** Trough-to-peak time (b) and half amplitude width of the negative peak (c) provided separation of units into excitatory and inhibitory neurons with typical spike shapes. **(C)** Examples of regular spiking excitatory neuron (top) and fast spiking inhibitory neuron (bottom) as recorded as recorded in an experiment. **(D)** Cross correlograms of neuron-pairs that indicate excitatory (left) and inhibitory (right) monosynaptic connections as an example. Horizontal lines of “+” symbols indicate high and low global confidence intervals obtained from spike-jittered average. **(E)** Spike trains of sorted units (unclassified) in wakefulness (0%) and anesthesia (6%), as an example. Spike patterns are sparser and more fragmented during anesthesia as compared to wakefulness.

## Results

Simultaneous extracellular recording and electrical stimulation was performed in nine rats using 32-site microelectrode arrays in primary visual cortex at three levels of anesthesia and wakefulness. **Figure 1E** presents an example of the typical firing pattern of sorted single units in wakefulness and anesthesia prior to microstimulation. The regular firing pattern in wakefulness converts to a typical intermittent firing pattern of the same neurons in anesthesia.

Microstimulation was applied to each electrode site was then stimulated one by one in random order at 2Hz repetition rate using charge-balanced biphasic pulses. We compared the effect of stimuli with different phase duration, amplitude, and temporal order on the firing response of neurons. In all stimuli, the maximum current was 40 μA with 60 μs pulse width and balanced by another phase with opposite polarity as illustrated in **Figure 2**. Putative excitatory and inhibitory neurons were not distinguished in this analysis. Cathodic-first charge-balanced stimuli activated more neurons than anodic-first charge-balanced stimuli, although this difference was small. All subsequent experiments were performed with asymmetric biphasic pulses with 40 μA maximum intensity and 60 μs pulse width, balanced with 8 μA intensity and 300 μs pulse width.

**Figure 2.**
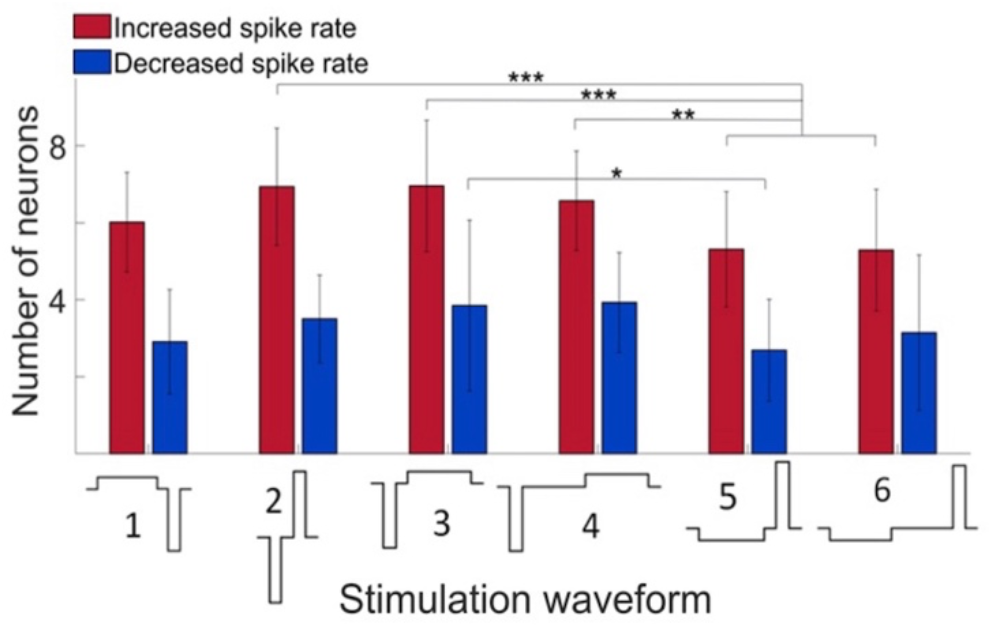
Effect of stimulus waveform on the number of neurons showing significant change in firing rate. Maximum stimulus current was 40 μA with 60 μs pulse width and charge balanced. Responses elicited from various stimulation sites were pooled. All data were obtained in wakefulness. Data are from 4 animals. Mean ±SD, *: p<0.05, **: p<0.01, ***: p<0.001.

**Figure 3** shows the time course of spike responses of putative excitatory and inhibitory neurons. Neurons were classified by their spike waveform features. Microstimulation generally caused an early increase, within 10 ms, followed by a prolonged (11-100ms) decrease in neuron firing. Both excitatory and inhibitory neurons showed this behavior. Neurons responding to stimulation were scattered throughout the electrode array. Despite the smaller number of inhibitory neurons recorded, their total spike count was higher due to their generally higher average spike rate. The maximum spike rates attained post stimulus at 6, 4, 2, and 0% desflurane were 2.3, 4.0, 4.5, and 6.5 spikes/s, respectively, for excitatory neurons and 10.0, 15.9, 18.4, and 25.0 spikes/s, respectively, for inhibitory neurons.

**Figure 3.**
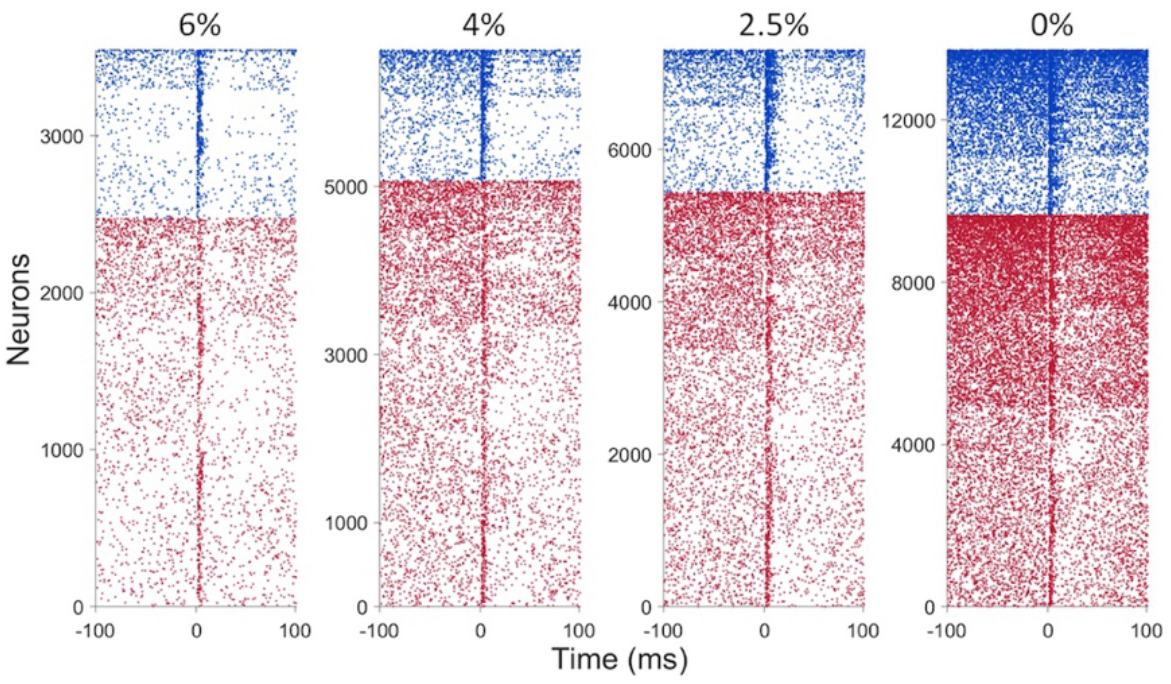
Spike raster plots of the effect of microstimulation on putative inhibitory (blue) and excitatory (red) neuron firing in four conditions at inhaled concentration of the anesthetic desflurane indicated on top. Each line corresponds to a neuron responding to a different trial, i.e., to stimulation at a different electrode site. Neurons with at least one spike within ±100 ms of stimulation only were included. Neurons of both types were sorted vertically by their firing rate. Stimulus is at time 0, charge-balanced, biphasic, asymmetric, cathodic-leading, pulses with 40 μA maximum intensity, 60-60-300 μs durations, delivered at 2Hz. Data are from 9 rats.

**Figure 4A** compares the change in spike rate of each neuron as a function of distance from the stimulation site separately for the early (1-10ms) and late (11-100ms) response. The magnitude of spike rate increase decayed with the neuron’s distance from the stimulation site. In contrast, the magnitude of spike rate decrease was essentially independent of the distance from the stimulation site. This pattern was essentially conserved across all conditions. The number of neurons responding with a significant increase or decrease in spike rate to stimulation in the two epochs is illustrated in **Figure 4B**. The magnitudes of the early increase and late decrease tended to increase slightly at lighter levels of anesthesia and upon waking although these effects were not statistically significant. Approximately twice as many neurons decreased than increased their spike rate following microstimulation (p<0.0001).

**Figure 4.**
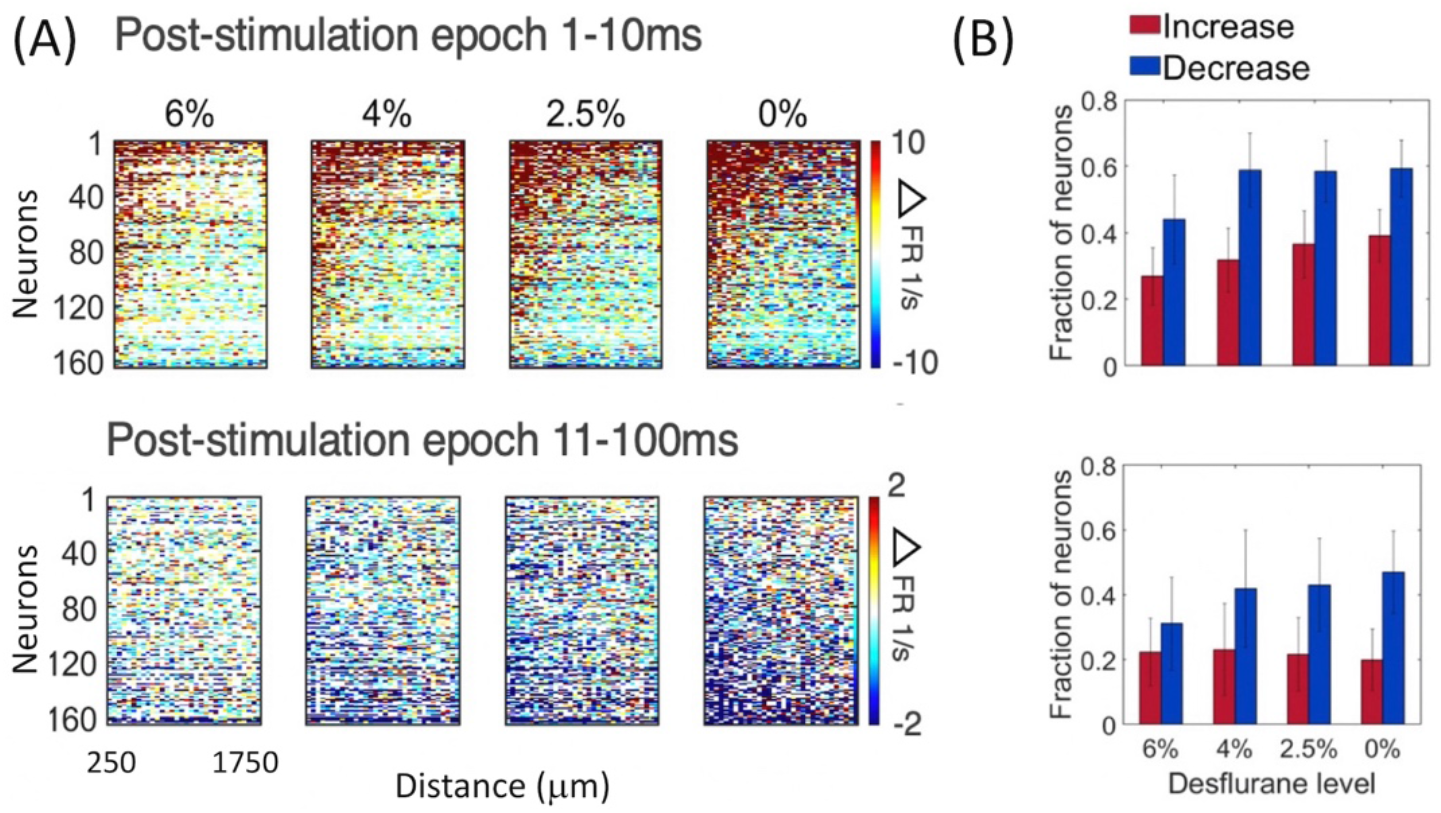
Change in firing rate as a function of distance from stimulation site in four conditions and two post-stimulus epochs. **(A)** Change in spike rate change of individual neurons. For each neuron, the 32 stimulation channels are arranged left to right in the order of increasing distance from stimulation site. Neurons are arranged top down from maximum increase (red) to maximum decrease (blue) of spike rate during the wakefulness (0%) and the same order is kept for the other three conditions. Desflurane concentrations are shown on top. Data are from 9 rats. **(B)** Fraction of neurons that produced statistically significant increase (red) or decrease (blue) in firing rate. More neurons showed decrease than increase of firing rate with microstimulation (p<0.0001), a difference qualitatively conserved across states. Stimuli were asymmetric, cathodic-leading, biphasic pulses with 40 μA maximum intensity, 60-60-300 μs durations, delivered at 2Hz.

We examined more closely how the early response of excitatory vs. inhibitory neurons varied as a function their distance from the stimulation site. This analysis was limited to the first 4 ms compatible with putative monosynaptic effects. As **Figure 5** shows, the number of excitatory neurons that significantly increased their firing rate decayed rapidly with distance from the stimulation site. This dependence was absent for inhibitory neurons suggesting that they were stimulated in a more distributed manner.

**Figure 5.**
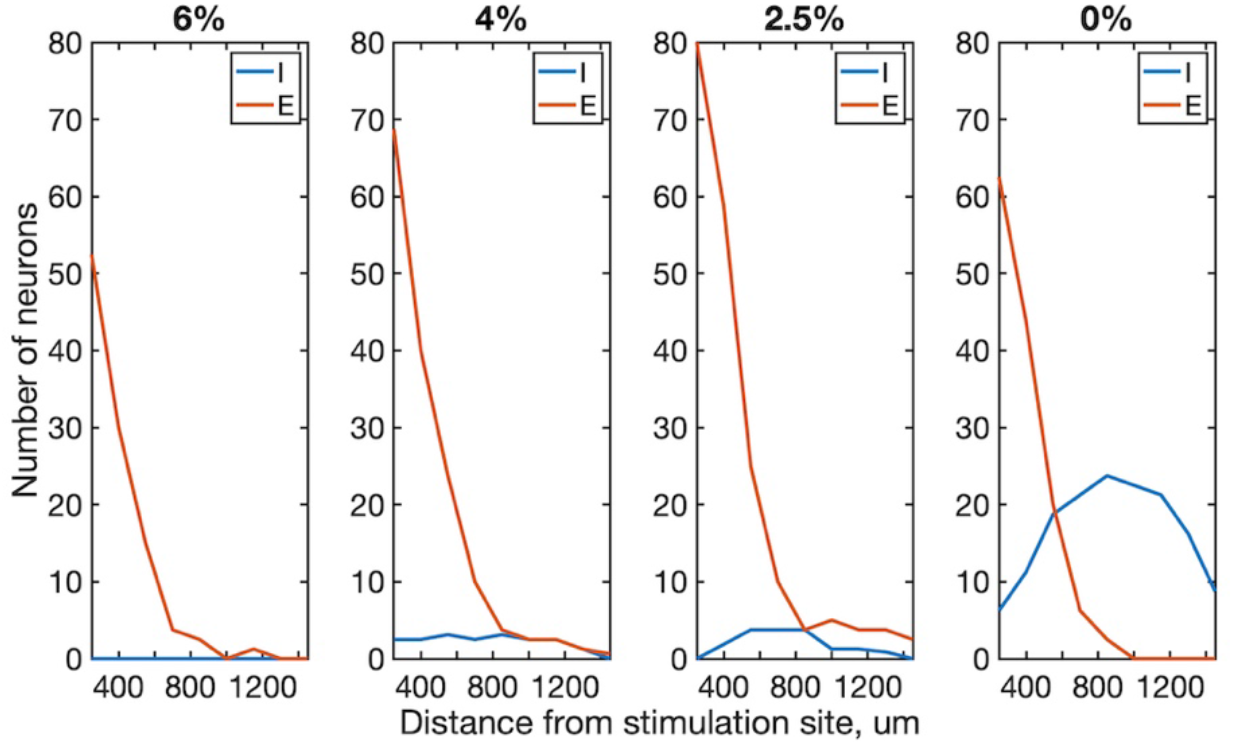
Effect of distance from stimulation site on excitatory and inhibitory neuron spiking within 1-4 ms post-stimulus interval in four conditions. The number of neurons whose firing rate exceeded 99% confidence interval of their prestimulus firing rate are plotted. Trials from all stimulus sites were combined. Data are from 4 rats. Desflurane concentrations is indicated on top. Stimuli were charge-balanced, asymmetric, biphasic, cathodic-leading pulses with 40 μA maximum intensity, 60-60-300 μs durations, delivered at 2Hz.

We then examined the spatial pattern of stimulus-induced (effective) connectivity of neurons. This was done in two different ways. First, a directed graph was constructed from the vectors from each stimulation site to all neurons whose firing rate was either significantly increased or decreased by microstimulation (**Figure 6A**). The resulting connectivity map illustrates the overall influence of microstimulation on the spiking of all recorded neurons. Comparing different conditions, effective connectivity was sparse under anesthesia, suggesting reduced overall neuronal excitability. The total number of positive effects (neurons with increasing spike rates) was significantly greater as the anesthetic was withdrawn. Thus, the effective connectivity map indicated progressively stronger influence of stimulation at lighter levels of anesthesia and especially in wakefulness. Moreover, negative effects (neurons with decreasing spike rates) were almost absent at all levels of anesthesia. Second, we estimated effective connectivity from putative pair-wise monosynaptic connections as determined from spike cross-correlograms separately for both pre- and post-stimulus periods. Identifying all monosynaptic connections and combining those in the same map, we derived a directed graph representing putative polysynaptic connectivity (**Figure 6B**). As with the previous method, higher connectivity was evident in light anesthesia and wakefulness and more in the post-stimulus than pre-stimulus phase.

**Figure 6.**
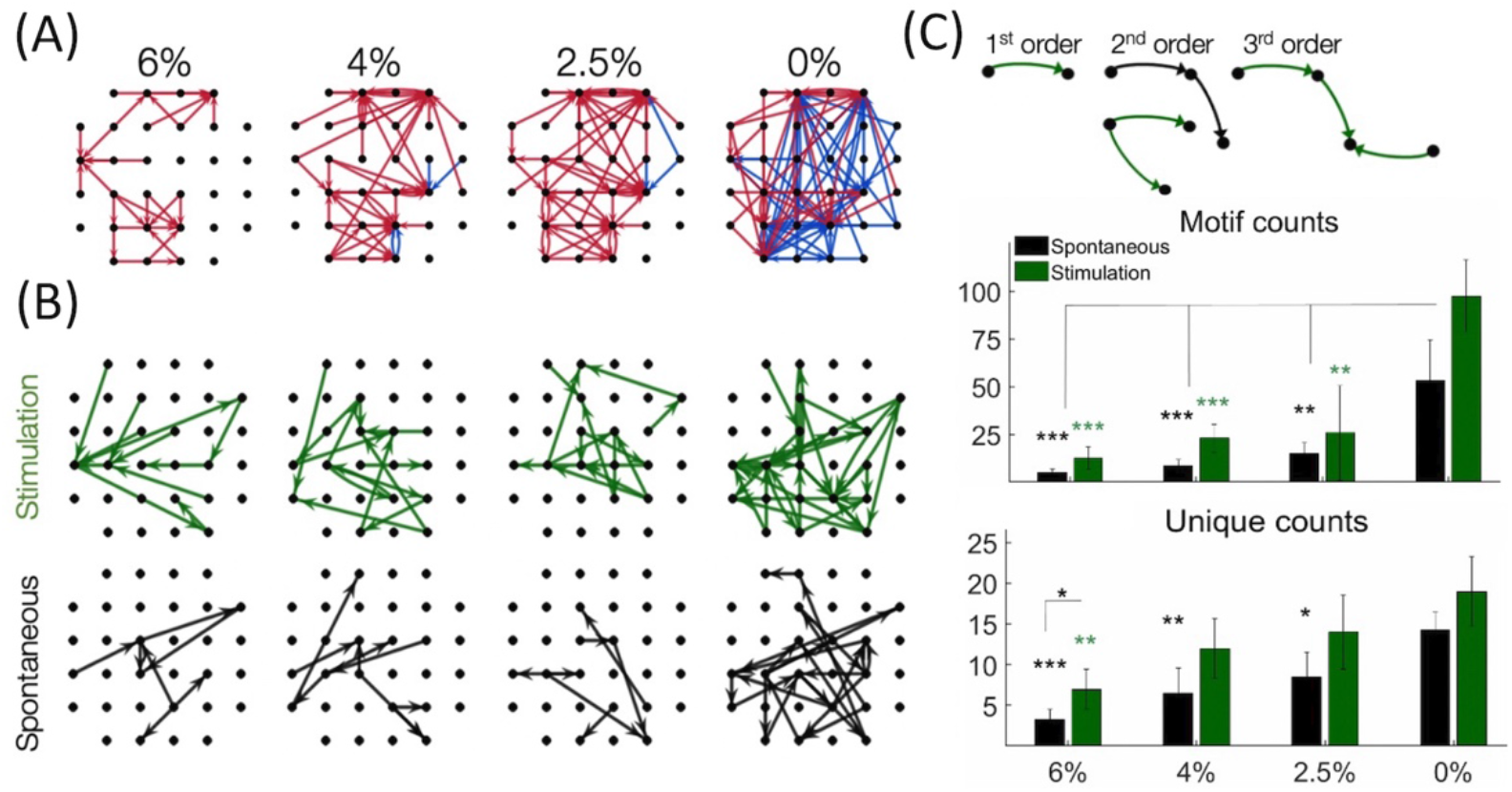
Effective connectivity of neurons determined by microsimulation. (**A**) Connectivity maps in one rat as an example. Arrows point from the site of stimulation to the site of significantly increased (red) or decreased (blue) spike rate within 1-4 ms post-stimulation (desflurane concentrations indicated on top). Desflurane concentrations is indicated on top. (**B)** Putative monosynaptic effective connectivity maps inferred from pairwise spike cross correlograms during stimulation. **(C)** Effect of anesthesia on the number of connectivity motifs as inferred from pairwise spike cross-correlograms before and after microstimulation. Examples of motifs of different order are illustrated on top. The counts of 1st, 2nd, and 3rd order motifs were combined. Unique counts reflect the diversity of distinct motif types. Desflurane concentration is on the horizontal axis. Data are from 4 rats. Mean ±SD, *: p<0.05; **: p<0.01; ***: p<0.001. Stimuli were charge-balanced, asymmetric, biphasic, cathodic-leading pulses with 40 μA maximum intensity, 60-60-300 μs durations, delivered at 2Hz.

To further examine the structure of the effective connectivity map, we identified low-level functional motifs of effective connectivity. A motif is a subset of patterns of putative monosynaptic connections arranged in series or in diverging or converging patterns, leading to multiple variants of second, third and higher order motifs (Recanatesi et al., 2019). First order motifs consist of a single monosynaptic connection; second order motifs contain two connections; and third order motifs contain three connections. Second and third order motifs have multiple variants defined by their connectivity profile. **Figure 6C** shows that microstimulation increased the total number of detected motifs, thus enhancing monosynaptic and polysynaptic connectivity at all levels of anesthesia although this effect did not reach significance at 0% desflurane in this comparison (for statistics see figure). Similar results were obtained for the number of unique motifs, suggesting that their diversity increased after microstimulation. In addition, both the total number of motifs and their diversity significantly increased as the anesthetic was withdrawn and reached maximum in wakefulness (see figure for statistics). **Figure 7** compares the change in the number of motifs of different orders separately. A progressive increase in the number of all motifs with decreasing anesthetic concentration was evident both pre-stimulus and post-stimulus (p<0.0001, ANOVA), except in post-stimulus first order motifsIn addition, microstimulation increased the number of motifs of all orders (p<0.005, ANOVA) with the most dramatic effect seen in motif order 3 (for pairwise statistical comparison see figure).

**Figure 7.**
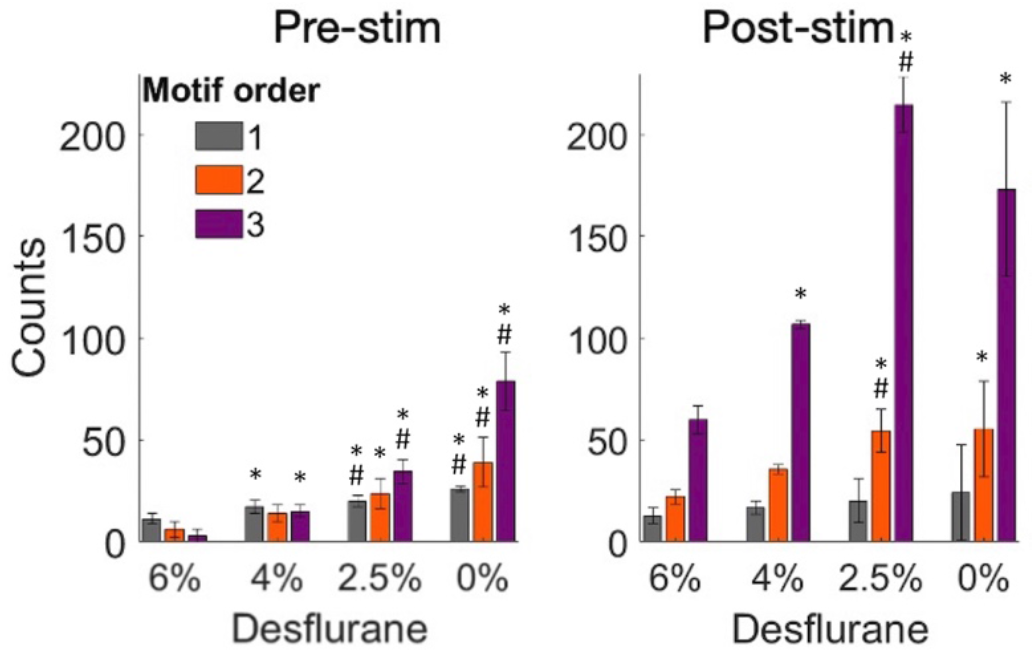
Change in the number of motifs of orders 1-3 as a function of experimental condition before and after microstimulation. Motifs were derived from pairwise spike cross-correlograms. Stimuli were charge-balanced, asymmetric, biphasic, cathodic-leading pulses with 40 μA maximum intensity, 60-60-300 μs durations, delivered at 2Hz. *: p<0.05 vs. at 6%, #: p<0.05 greater than at next higher desflurane concentration.

## Discussion

The main goal of this work was to apply intracortical microstimulation to map effective connectivity (Aertsen et al., 1989) of a local neuronal circuit and examine how anesthesia may alter such connectivity in a dose-dependent manner. Microstimulation was chosen as an effective means to modulate local neuronal activity and its propagation across adjacent and remote recording sites (Kwan and Dan, 2012;Kumar et al., 2013). Probing local circuitry by causal perturbation provides additional insight into neuronal connectivity underlying cortical information processing over that obtainable by recording spontaneous or physiological stimulus-related activity (Clark et al., 2011;Kumar et al., 2013;Cicmil and Krug, 2015;Sadeh and Clopath, 2020). Further, by investigating the concentration-dependent effect of anesthesia on intracortical stimulus-related connectivity has a unique potential to help better understand how neuronal networks function in altered states across experimentally controlled levels of consciousness (Arena et al., 2021).

Consistent with prior studies we found that intracortical microstimulation generally produced a biphasic neuronal response consisting of an early increase in firing within 10 ms followed by a transient suppression of 100 ms or so (Butovas and Schwarz, 2003;Sombeck et al., 2022;Yun et al., 2023). This biphasic pattern was conserved at all levels of anesthesia although the spike rate increases were lower during anesthesia than in the awake state. The exact duration of these phases was reported to vary with stimulus intensity (Yun et al., 2023), although this aspect was not investigated further here.

Also, as found before (McIntyre and Grill, 2000;Voigt and Kral, 2019), cathodic-leading charge-balanced stimulus waveforms were more effective in stimulating neuron firing than the reverse waveforms, although this difference in our preparation was quite small. Experimental and computational studies suggested that local cells are preferentially activated by cathodic-first asymmetrical charge-balanced biphasic stimulus waveforms, while fibers of passage is affected more by anodic-phase-first asymmetrical charge-balanced biphasic stimulus waveforms (Cogan, 2008;Voigt and Kral, 2019). However, it was also shown that stimulus polarity and asymmetry influence the probability and localization of neuron activation in an opposite manner (Stieger et al., 2022). Also, computational modeling suggests that the initial mechanism of activation is an antidromic propagation to the soma following axonal activation (Kumaravelu et al., 2022). Synaptic transmission could then mediate the further propagation of activation.

To better understand the synaptic involvement in the early stimulus response, we examined putative monosynaptic connectivity estimated from post-stimulus spike cross-correlograms within 1-4 ms. Transsynaptic responses to single-pulse cortical electrical stimulation were reported in a comparable time frame (Sombeck et al., 2022). We found that under anesthesia, very few inhibitory neurons showed monosynaptic response, which however was increased substantially in wakefulness, suggesting a normalization of E/I balance. Moreover, the wide spatial distribution of stimulus-responding inhibitory neurons implied their excitation by horizontal axonal projections that extend over 1mm (Kisvarday, 1992). Prolonged suppression of firing rate could be mediated by a network of electrically coupled inhibitory interneurons (Butovas et al., 2006) that are particularly dense around the stimulation electrode (Overstreet et al., 2013). This does not seem to be the case for excitatory neurons, however. Indeed, the decay of the number of excitatory neurons with distance from the stimulation site could be related to their relatively sparse connectivity (Overstreet et al., 2013). A spatial decay over 1mm similar to that in our data was previously reported (Eles et al., 2021). Alternatively, the decay may be due to electrotonic excitation via field potentials that decay exponentially with distance from the stimulation site.

The main finding of our study was a graded reduction in cortical effective connectivity at increasing depth of anesthesia. This was consistent with an overall reduction in neuronal firing rate impeding ongoing neuronal interactions and the propagation of activity to remote sites. Moreover, negative effects of microstimulation (neurons with decreasing spike rates) were virtually absent in anesthesia suggesting that they were at the minimum baseline activity. This is consistent with the direct effect of volatile anesthetics on cortical firing by augmenting inhibitory neurotransmission (Hentschke et al., 2005). The directed effective connectivity graphs derived from functional monosynaptic connections confirmed these findings revealing sparser connectivity in anesthesia compared to the dense connectivity in wakefulness that was further augmented in the post-stimulus phase.

To gain further insight into network connectivity, we also analyzed the prevalence of simple network motifs in different conditions. Such motifs have been considered as building blocks of complex recurrent networks. The motifs’ statistical prevalence can determine overall network properties, such as dimensionality, code compression, computational flexibility, network input-output response, and memory (Hu et al., 2014;Recanatesi et al., 2019). In the rat visual cortex, recordings of layer 5 pyramidal neurons revealed the presence of small, strongly connected network units that determine overall network connectivity (Song et al., 2005). Our analysis revealed a sharp difference in the overall number of motifs between wakefulness and anesthesia consistent with the rapid collapse of network connectivity. This difference was most expressed in the number of 3^rd^ order motifs in the pre-stimulus condition. Although we were not able to identify motifs of higher than 3^rd^ order due to the limited size of the electrode array, it could be surmised by extrapolation that higher order motifs would be even more affected. The additional finding that microstimulation did not significantly increase the number of motifs in the awake condition could mean that they were already at their near maximum density.

A few limitations of the current study are recognized. First, due to the sparse sampling of neuronal network, effective connectivity maps represent a small subset of true neuronal connectivity. A complete reconstruction of the neuronal network would theoretically require stimulating all single neurons and their combinations of the entire neuron network (O’Doherty et al., 2012;Kumar et al., 2013;Sadeh and Clopath, 2020). Nevertheless, the observed change in graph density in our sample should reflect the general trend of the anesthetic effect. Second, neuronal connectivity as determined here could be in part from synaptic transmission and in part from electrical fields through a mixture of antidromic and orthodromic activation (Butovas and Schwarz, 2003). In our experiments, the probability of evoking excitatory responses decayed with distance rapidly, which could be due to the increasing sparsity of synaptic terminals reached or the weakening of electrical field with distance. Although cross-correlograms with short time-lag likely reflect monosynaptic spike transmission probability between pairs of neurons, they can be confounded by common inputs with comparable temporal delay. Thus, the neuron connections derived by either method applied here are approximate and connections should be considered “putative”. Third, extracellular microstimulation affects not a single cell but an unknown population of cells and passing axons surrounding the stimulation electrode. These difficulties could be mitigated by neuronal redundancy, i.e., that many neighboring neurons often respond to similar stimulus features, that a relatively small group of cells can drive network activity in similar fashion (Yassin et al., 2010;Kwan and Dan, 2012;Emiliani et al., 2015;Bernardi et al., 2021), and that frequently recurring neuronal firing sequences represent a small fraction of all possible patterns (Luczak et al., 2007;Luczak et al., 2009;Luczak and Maclean, 2012). Thus, effective connectivity as derived here is to be interpreted as the influence of a local neuronal population on individual neurons at a distance. Experiments and computer simulations suggest that such an approach to measure neuronal “embeddedness” is a valid approach for network discovery (Vlachos et al., 2012;Kumar et al., 2013). In addition, the 2^nd^ and 3^rd^ order network motifs could be more correctly interpreted as neuron-to-neuron connectivity, especially when assessed through a physiological method such as the spike cross-cross correlogram. Fourth, one could contend that the observed changes in connectivity may in part be accounted for by the anesthetic-induced decrease in overall firing rate. However, this is not thought to be the primary contributing factor of effective connectivity changes because stimulus-induced neuronal firing is often significantly elevated during anesthesia, especially at deep anesthetic levels (Hudetz and Imas, 2007). Finally, trial-to-trial variability due to ongoing activity limits the accuracy at which an invariant network can be reconstructed. Nevertheless, it is believed that the microstimulation-based perturbational approach augments the reproducibility of the mapped network by transiently suppressing background activity (Cheney et al., 2013). In the future, optical methods that allow more precise control of the number and type of directly stimulated neurons (Cicmil and Krug, 2015;Cavelli et al., 2023) including optogenetic stimulation with micro-LEDs (Buzsaki et al., 2015;Wu et al., 2015;Kim et al., 2020) could be used. In addition, distinct temporal patterns of stimulation pulses could also be used to control the affected neuronal population (Eles et al., 2021) and one could also target the stimulation to specific cortical layers (Voigt et al., 2017;Urdaneta et al., 2021).

In summary, the present work demonstrates the anesthetic modulation of effective connectivity in rat visual cortex as probed by microstimulation *in vivo*. The suppression of network connectivity was associated with the preferential loss of higher order network motifs. The results are relevant to understanding the mechanisms of anesthetic-induced impairment of cortical information processing and by inference, loss of consciousness.

## Acknowledgements

Research reported in this publication was supported by the National Institute of General Medical Sciences of the National Institutes of Health under award number R01-GM056398 and the Center for Consciousness Science, Department of Anesthesiology, University of Michigan Medical School, Ann Arbor, Michigan, USA. The content is solely the responsibility of the authors and does not necessarily represent the official views of the National Institutes of Health. The authors thank the assistance of Dr. Shiyong Wang in conducting the experiments, Jake Schut in data analysis, and Kathy Zelenock in laboratory operations.

